# An optimized high-throughput colorimetric assay for phytic acid quantification

**DOI:** 10.1101/2025.07.30.667628

**Authors:** Ahmed O. Warsame

## Abstract

Phytic acid (PA) is the primary storage form of phosphorus in seeds and is considered an anti-nutritional factor due to its ability to chelate essential minerals, thereby reducing their bioavailability. However, identifying low-PA mutants in large populations requires a cost-effective, accurate, and high-throughput screening method. Gao *et al*. (2007) developed a colorimetric method that is cost-effective and accurate for quantifying PA in soybeans. However, the throughput of this method was relatively low. In this study, we modified some critical steps of the original protocol to improve throughput. The accuracy and reproducibility of the modified protocol were validated by comparison with those of the original method and a commercially available PA quantification kit. The new high-throughput protocol showed high reproducibility and successfully distinguished existing low-PA mutants from their wild-type parent. The protocol was then used to screen 202 accessions from a pea diversity panel, which revealed wide genetic variation in PA content. We identified two novel low-PA accessions, JI0383 and JI3253, with 69% and 48% reductions in PA, respectively, compared to the population mean. This cost-effective method is expected to help researchers and breeders accelerate the development of low-PA crops to meet the current demands for high-quality plant-based foods.

## INTRODUCTION

Phytic acid (PA), or myo-inositol hexakisphosphate (InsP6), is the primary storage form of phosphorus in seeds, accounting for up to 80% of total seed phosphorus (Bohn et al., 2008). PA plays an important role in plant physiology, particularly in seed germination, seedling development, and stress responses (Pramitha et al., 2021, Sparvoli and Cominelli, 2015). However, PA is considered an anti-nutritional factor in human and animal diets because of its ability to chelate essential minerals, such as iron and zinc, thereby reducing their bioavailability (Moore et al., 2018, Bangar et al., 2017, Sparvoli and Cominelli, 2015). Considering that many staple crops contain relatively high concentrations of PA (Wang et al., 2022, Pramitha et al., 2021), this exacerbates the global prevalence of micronutrient malnutrition in population groups that depend on plant-based diets. Additionally, in high-income countries, shifting towards environmentally sustainable food systems requires increasing the consumption of pulses (Willett et al., 2019), which are naturally PA-rich and potentially lead to mineral deficiencies in specific demographic groups, notably women of reproductive age (Beal et al., 2023). Therefore, lowering PA concentration is necessary to unlock the nutritional and environmental benefits of pulse crops.

From a nutritional perspective, foods should ideally contain low PA and high mineral density, particularly iron and zinc. However, achievements in developing staple legumes with high mineral contents have yielded only modest success (Bouis and Saltzman, 2017). One major challenge is that excessive mineral accumulation can trigger toxic responses in plants, making mineral levels tightly regulated (Harrington et al., 2024). Thus, a practical breeding strategy could be to introduce low-PA alleles into cultivars that already possess relatively higher mineral content.

Screening large numbers of accessions in natural diversity or mutant collections requires an accurate and efficient method for quantifying PA content. Several methods have been described for PA quantification including anion exchange chromatography (AEC), high-performance liquid chromatography (HPLC), and nuclear magnetic resonance (NMR) spectroscopy (Gao et al., 2007). A major advantage of these methods is their ability to distinguish between InsP6 (PA) and lower inositol phosphates (≤ InsP5) (Marolt and Kolar, 2020). However, these methods are expensive, require specialised equipment, and are unsuitable for high-throughput analyses. In contrast, cost-effective colorimetric methods that are amenable to higher throughput have been developed. One such method is the molybdate blue assay, which is based on the reaction between phosphate and molybdate to form a blue-colored complex. Using this principle, the commercially available Megazyme kit uses a combination of phytase enzymes to quantify total phosphate and PA-bound phosphate, and is widely used for PA quantification (Kumar et al., 2021, Gyani et al., 2020, Cominelli et al., 2018). However, this method is relatively expensive and involves time-consuming sample handling steps. The molybdate blue assay has also been used to quantify the concentration of inorganic phosphate (P_i_), and has facilitated the identification of several low-PA mutants in different crops (Warkentin et al., 2012, Qamar et al., 2024, Gyani et al., 2020). Because this method assumes that samples with high P_i_ are potential mutants with impaired PA biosynthesis, it may not detect mutants with reduced overall seed phosphate content caused by mutations affecting phosphate transport from lower tissues to the developing seeds. For instance, Yamaji et al. (2017)Such rice mutants have shown significantly lower P_i_ content in grains than the wild type. Therefore, the P_i_-based molybdate blue assay is insufficient to detect such valuable mutants. A third colorimetric method that directly measures PA is the Wade reagent assay (Vaintraub and Lapteva, 1988) which was further modified by Gao et al. (2007). This method exploits the high affinity of PA for Fe3^+^, where the pink colour of the Fe^3+^–sulfosalicylate complex (Wade reagent) is depleted in proportion to the concentration of PA in the test sample. This offers a simpler and more cost-effective alternative that can be used for both quantitative and qualitative analyses of PA content. The method has been used for different crop species, including rice (Panda et al., 2023), wheat (Williams et al., 2024), chickpea (Basu et al., 2019), mung bean (Dhole and Reddy, 2016), and groundnut (Hande et al., 2013). However, the utility of this method for large-scale screening studies is limited by its low throughput (Wen et al., 2022). Therefore, to improve throughput, this study aimed to further modify and refine the colorimetric protocol proposed by Gao et al. (2007). All the steps of the modified high-throughput PA assay were performed in 96-well plates. This improved high-throughput protocol was highly reproducible and allowed the identification of novel low-PA pea accessions.

## MATERIALS AND METHODS

Pea (*Pisum sativum* L) accessions were obtained from the John Innes Centre Germplasm Resource Unit (website). The PCGIN diversity panel (https://pcgin.org/) consists of 230 lines covering the diversity spectra of the pea germplasm held at the John Innes Centre (Jing et al., 2010). In addition, two low-PA mutant pea lines (1-150-81 and 1-2347-144) and their parental line (CDC Bronco) were kindly provided by Professor Tom Warkentin of the University of Saskatchewan (Warkentin et al., 2012). The lines were grown in field plots at the Dorothea de Winton Field Station in Norwich in 2021. The final number of accessions for PA analysis was 202.

### Sample Preparation

Approximately 10 g of dry pea seeds were ground using an MMT 40.1 Multi-use milling tube (IKA, Staufen, Germany) for 1.5 minutes at 25,000 rpm. To ensure homogeneity, the samples were then sieved through a 0.250 mm mesh fitted to a custom-made frame. Subsequently, the samples were dried overnight at 70 °C.

### Chemical reagents

All reagents utilized in this study were of analytical grade and were sourced as follows: the PA/Total Phosphorus Kit (CAT#: 700004327) was obtained from Megazyme Inc.; Phytic Acid Sodium Salt Hydrate (CA#: P8810), Ascorbic Acid (CAT#: A7506), Sulfuric Acid (CAT#: 258105), Ammonium Molybdate Tetrahydrate (CAT#: 09878), NaCl (CAT#: S9888), and Trichloroacetic Acid (CAT#: T8657) were sourced from Sigma-Aldrich; Sulfosalicylic Acid (CAT#: 11475533), Nitric Acid (CAT# 10149982), Hydrogen Peroxide (Fisher Scientific, CAT# 10530341), and HCl (CAT#: 10316380) were acquired from Fisher Scientific; and Iron(III) Chloride Hexahydrate (CAT#: 31232) was obtained from Fluka.

### PA quantification

#### Total phosphorous method

The principles underlying the method are described in McKie and McCleary (2019), and the reagents are available as a commercial kit (K-PHYT) from Megazyme (Bray, Ireland). The assay was conducted in accordance with the manufacturer’s protocol (Megazyme Inc., Bray, Ireland). Briefly, 1 g of pea seed flour was weighed into 50 ml Falcon tubes and digested with 20 ml of 0.66 M HCl overnight. Subsequently, a 1 ml aliquot was transferred to a 1.5 ml tube and centrifuged at 15,000 × g for 10 minutes. Thereafter, 0.5 ml of the supernatant was neutralized with 0.5 ml of 0.75 M sodium hydroxide solution. In parallel steps, free phosphorus and total phosphorus were quantified, the latter being released through a two-step enzymatic dephosphorylation process. The molybdenum blue method was employed to determine phosphorus concentration, utilizing a standard calibration curve ranging from 0.125 µg/ml to 8 µg/ml. The colour intensity of the samples was measured at 655 nm using a microplate reader (ClarioStar, BMG LabTech).

#### Gao et al. 2007 method

This was performed following the original protocol by Gao et al. (2007). Briefly, ∼500 mg seed flour was mixed with 14 ml of 0.64 M HCl overnight. Tubes were centrifuged and the supernatants were into new tubes containing 1 g of NaCl to remove matrix components that could interfere with the colorimetric reaction (Gao et al., 2007). After incubating on ice for 1 hours, samples were centrifuged and the clear supernatants were further diluted and used for analysis. The concentration of PA was determined using Wade Reagent, which was prepared by dissolving 300 mg sulfosalicylic acid and 30 mg Ferric Chloride Hexahydrate in 100 mL of Milli-Q water. For colour development, 3 ml sample was mixed with 1 ml Wade Reagent, PA content was calculated from seven standard curve samples ranging between 20 to 80 µg/ml.

#### Modified high-throughput PA method

The assay was conducted using 96-well plates rather than tubes (refer to the detailed protocol in Figure 1). Approximately 60 mg of seed flour was weighed into 1.2-ml 8-strip tubes arranged in a 96-well format (QIAGEN, CAT# 19560). Each plate comprised 88 pea samples, including duplicates of 1-150-81 and CDC Bronco, which are a known low-PA mutant and its parental line, respectively. The remaining wells were allocated for the standard curve and blank samples in subsequent steps of the assay.

**Figure 1.**
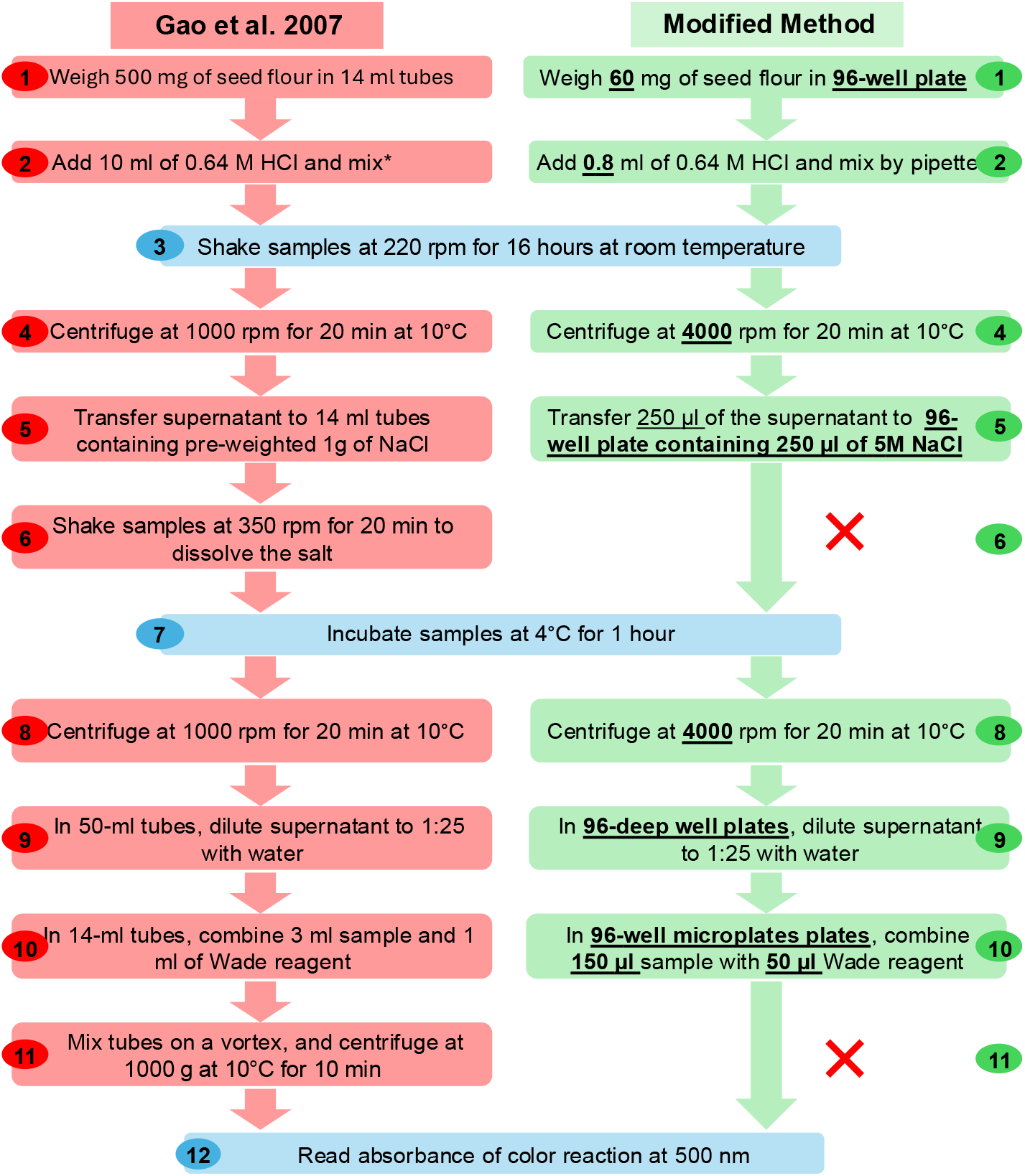
A step-by-step comparison of the original protocol by Gao et al. (2007) and the modified high-throughput assay. In the modified method, omitted steps are indicated by red “X” while other modifications are shown in bold text. The asterisk in step 2 indicates that Gao et al. (2007) did not specify how the samples were mixed; therefore, we mixed the samples by vortexing.

Using a 1 ml multichannel pipette, 800 µL of 0.64 M HCl was added to each sample. To mitigate clump formation, samples were gently mixed by pipetting while the acid solution was gradually added. Subsequently, the plates were sealed and positioned horizontally in beakers, which were then shaken overnight at 250 rpm at room temperature. The samples were centrifuged at 4000 rpm (∼2,720 × g) at 10°C for 20 minutes. Thereafter, 250 µL of the supernatant was transferred to new 1.2-ml deep-well plates, and 250 µL of 5 M NaCl solution was added. The samples were incubated at 4°C for 60 minutes, followed by centrifugation at 4000 rpm at 10°C for 20 minutes. Finally, 40 µL of the clear supernatant was transferred to new plates and diluted to a 1:25 ratio using 460 µL Milli-Q water. The colorimetric assay was performed in 96-well clear plates by combining 50 µL of Wade Reagent with 150 µL of the diluted samples, and the absorbance was measured at 500 nm.

### Elemental phosphorous analysis using ICP-OES

The concentration of total phosphorus was determined using Inductively Coupled Plasma-Optical Emission Spectroscopy (ICP-OES) as in (Harrington et al., 2023) with minor modifications. Approximately 100 mg of dried pea flour was digested with 1 ml of 68% nitric acid (Fisher Scientific, CAT# 10149982) and 250 µl of 30-32% hydrogen peroxide (Fisher Scientific, CAT# 10530341) at 95 °C for 16 h. The final acid concentration was reduced to 0.85% by diluting with Milli-Q water. All samples were analysed in duplicates and measured with three technical replicates. Each batch of samples contained a reference sample and pea sample, which were analysed in duplicate across all runs.

### Data analysis

Analysis of variance (ANOVA) was performed using R, and the significance of differences between means was determined using Tukey’s Honestly Significant Difference test (HSD) at p ≤ 0.05.

## RESULTS

To develop a high throughput assay for PA, the amount of flour from seeds was downscaled from 500 mg to 60 mg, and the extraction volume from 10 ml to 0.8 ml (Figure 1). However, the most important modification of the original protocol was to replace the solid NaCl with a 5M solution form. This eliminates the need to weigh equal amounts of salt into tubes and the time required to dissolve it in step 6 of the original protocol (Figure 1). In addition, because using 96-well plates is compatible with higher speed centrifuges, better separation of precipitates is achieved, which removes more sample matrix after salt treatment. Therefore, step 11 of sample mixing and centrifugation was omitted.

We validated the modified method by (i) comparing it with the original method and the commercial Megazyme kit, and (ii) its ability to distinguish high and low-PA samples using known low-PA mutants and their wild-type parental line. All three methods clearly distinguished the low-PA mutants (1-150-81 and 1-2347-144) from the wild-type CDC Bronco (Figure 2A). However, the PA content measured by the Megazyme kit was significantly lower than that quantified by Gao et al. (2007) and the modified version, which both showed 27% higher PA, on average. Conversely, except for the wild-type, the original and modified methods were not statistically different, and their higher values for PA content could be due to interference by unknown sample matrices, including P_i_ content. The Megazyme method measures only PA-Phosphate, whereas, according to Gao et al. (2007), the sample matrix, including its P_i_ content, may interfere with the Wade reagent. Nonetheless, the magnitude of the difference between the mutants and the wild-type remained nearly the same across the three methods, with 48-55% PA reduction in the mutants. In addition, the calibration curves were highly linear, with R^2^ values of 1 and 0.99 for the Megazyme kit (Figure 2B) and Wade reagent methods (Figure 2C), respectively. It is worth noting that the blank-corrected absorbance of the standards in the Wade reagent is negative because the intensity of the assay colour was inversely correlated with the concentration of PA.

**Figure 2.**
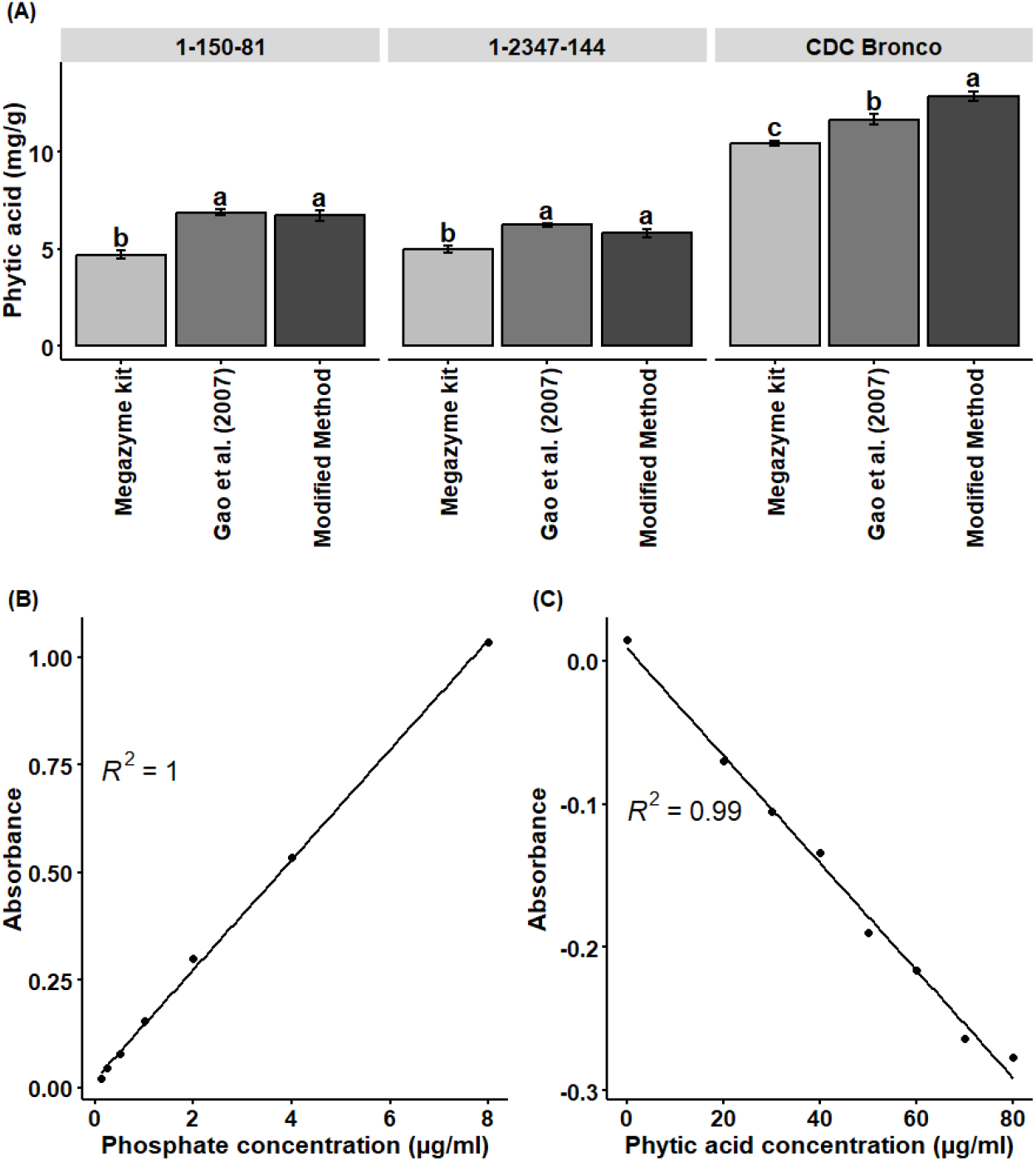
PA quantification using the Gao et al. (2007) protocol, the modified method, and the commercial Megazyme kit. (A) Comparison of PA content of low-PA mutants and their parental lines measured by the three protocols. The data represent the mean of three replicates, and the bars are standard errors. Tukey’s Honestly Significant Difference (HSD) mean comparison test was used, and bars with different letters indicate significant differences (p ≤ 0.05). (B) and (C) show the ranges and R2 for the phosphate standard used for the Megazyme kit and the PA standard used for Wade reagent assays, respectively.

To further test the utility of the modified protocol, we screened 202 accessions from the PCGIN pea diversity panel for PA content. There was wide genetic variation in PA content among the pea accessions, ranging from 5 to 21 mg/g, with an average of 16 mg/g (Figure 3A). The lowest PA was recorded in JI0383 and JI3253,with reductions of 69% and 48%, respectively, compared to the population mean. As expected, PA content had a strong positive correlation with the total phosphorous (P) content (Figure 3B). This high agreement between P measured by ICP-OES and the PA determined by the modified assay confirms that the majority of P in pea seeds is stored in PA and that the high-throughput protocol can accurately quantify PA in pea seeds.

**Figure 3.**
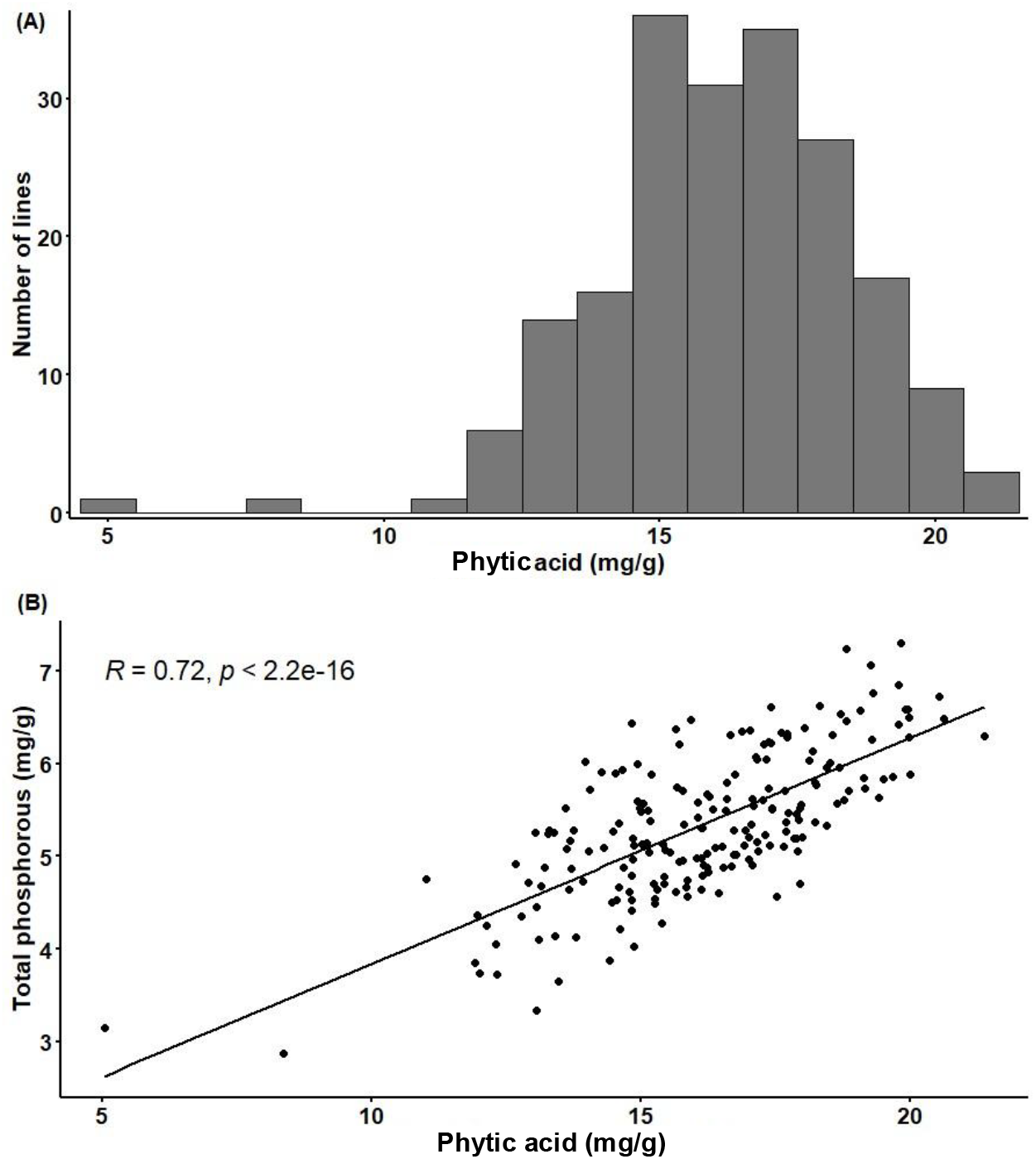
Variation in seed PA content among 200 pea accessions screened using high-throughput PA assay. Genetic diversity in pea for PA content (A) and its correlation with total phosphorous content measured by ICP-OES (B).

The reproducibility of the results obtained by the modified protocol was assessed by comparing PA content in two lines, 1-150-81 and CDC Bronco, which were replicated in six 96-well plates and measured in three batches on different dates (Figure 4A). There was no significant difference between the plates, except plates C and F for PA content of 1-150-81. In addition, the coefficient of variation (CV) of the PA content across the plates was 7.9% and 6.7% for 1-150-81 and CDC Bronco, respectively. This indicated the high reproducibility of the data across the plates and batches.

**Figure 4.**
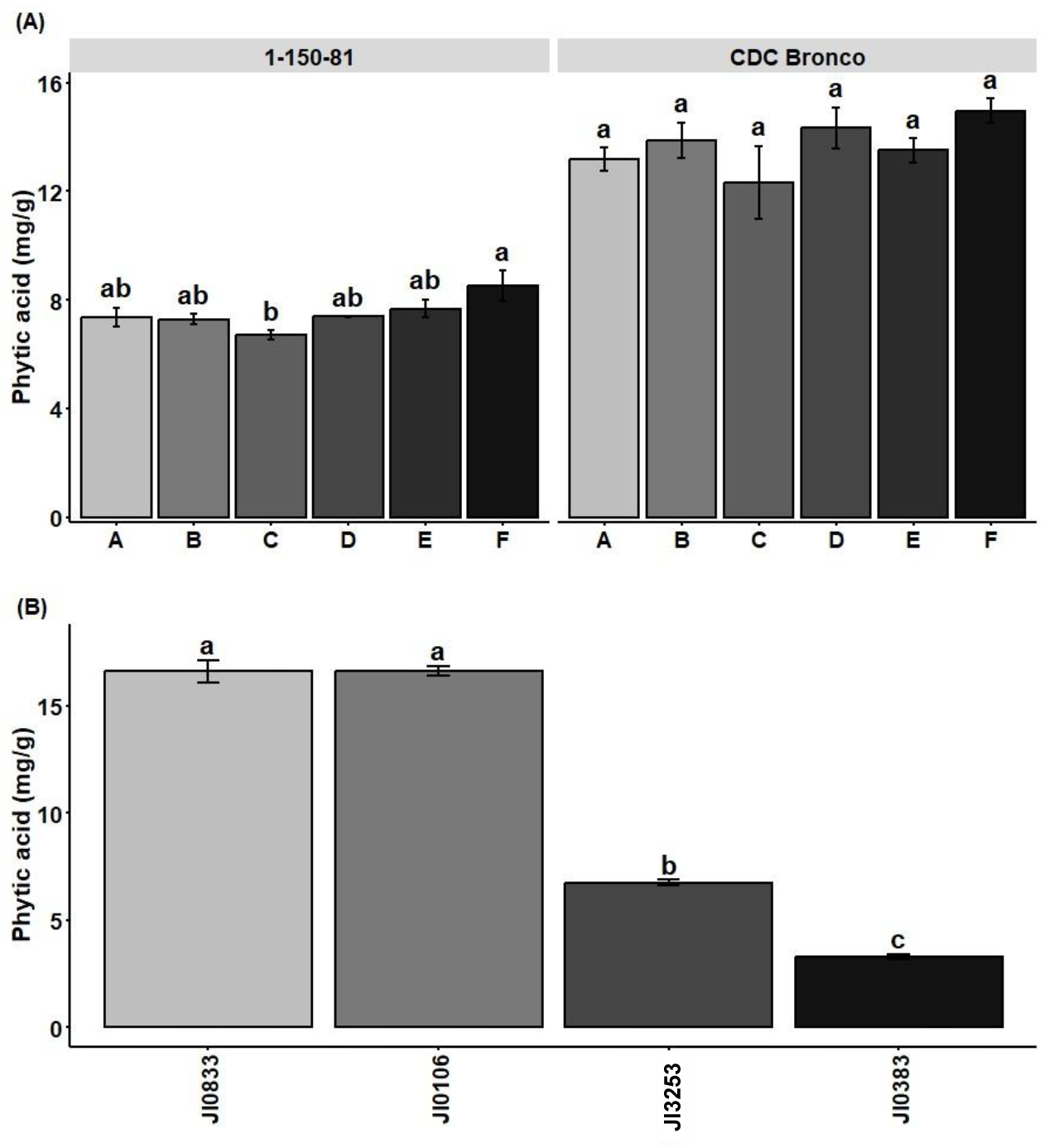
Reproducibility and accuracy of high-throughput assay. (A) PA content of a low-PA pea mutant line (1-150-81) and its parental line measured in different plates (A-F) and batches (batch1 = A&B, batch2 = C&D, batch3 = E&F). (B) The PA content of two lines in the pea diversity with very low AP, along two high-PA lines, were reanalysed using the Megazyme kit. Bars with different letters are significantly different (p ≤ 0.05) according to Tukey’s Honestly Significant Difference (HSD) mean comparison test.

Finally, to confirm the low PA content of JI0383 and JI3253, we reanalysed them along two high-PA accessions using the Megazyme kit (Figure 4B). This confirmed that these lines were novel genetic variants with very low PA content. In addition, the magnitude of the difference between the low- and high-PA lines was nearly the same in the Megazyme kit and the modified assay. For instance, compared with JI0833 and JI0106, the reduction in PA in JI0383 was 80% using the Megazyme kit and 77% using the high-throughput assay.

## DISCUSSION

Enhancing mineral bioavailability by lowering the phytic acid (PA) content in food crops can complement ongoing efforts to address the prevalence of malnutrition. Because of its nutritional significance and the growing demand for plant-based foods, this trait is expected to be a major breeding objective. Several low-PA lines have been developed for many crops, including pea (Warkentin et al., 2012), soybean (Punjabi et al., 2018, Gillman et al., 2009), common bean (Cominelli et al., 2018), rice (Yamaji et al., 2017, Tong et al., 2017, Qamar et al., 2024), and maize (Abhijith et al., 2020, Pilu et al., 2003). These materials are not widely available and are subject to IP restrictions, necessitating the development of local genetic materials, either from natural diversity or through mutagenesis programs. Such endeavours require screening of large populations, and the available methods for PA quantification are expensive and have low throughput. In this study, we have developed a refined version of the colorimetric method described by Gao et al. (2007). In the new protocol, we reduced the sample size and extraction volume by 10-fold and omitted or modified some steps to considerably improve the throughput. This modification is particularly valuable when working with limited seed quantities or when analysing large populations.

By comparing the modified assay with the original protocol and the commercial Megazyme kit, we showed that these modifications did not compromise the accuracy of the protocol. The protocol can screen 528 samples every two days (six plates), assuming that the samples were already weighed in 96-well plates, and one person handled the protocol. In the case of extremely large sample numbers, the throughput of the protocol can be further increased by grinding the seeds in 24, 48, or 96 batches using the Geno/Grinder® 2010 Tissue Homogeniser (see different options at https://cpsampleprep.com/). In addition, an electronic balance with the functionality to print data into an Excel file can enhance efficiency and minimise errors. In terms of the cost per sample, this method is extremely cheap and is a fraction of the cost of the Megazyme kit (50 assays).

Using the optimised protocol, we identified two pea accessions with very low PA contents. Based on their history, the two accessions are not related. The JI0383 accession is a garden pea line with wrinkled seeds and was donated by Elsoms Seeds Ltd. to the John Innes Centre in 1968. JI3253, also known as Cameor, is a French field pea cultivar with round seeds that has been extensively used in pea genetic studies, including the first full genome sequence (Kreplak et al., 2019) and genetic mapping populations (Ellis et al., 2023, Klein et al., 2020). Considering this breeding background and based on our own observations of the yield performance of these accessions both in the field and glasshouse, the loci conferring low-PA had no negative impact on their agronomic potential. This is crucial because of the agronomic penalties for PA reduction reported in pea (Warkentin et al., 2012), rice (Qamar et al., 2024, Zia ul et al., 2019), and maize (Abhijith et al., 2020). It appears that mutations in the PA biosynthesis pathway, which are often associated with the overaccumulation of P_i_ in the seed, have a negative impact on seed viability and germination. In contrast, Yamaji et al. (2017) showed that mutating the rice SULTR-like phosphorus distribution transporter (SPDT), which is expressed in the xylem region of nodes and controls the allocation of phosphorus to the grain, significantly reduces PA without compromising yield and seed germination. Our preliminary data (not shown) showed that these two low-PA pea lines accumulated a higher P_i_ content in their leaves, which is characteristic of impaired phosphate transport from lower vegetative tissues to the grains (Yamaji et al., 2017).

In conclusion, the new protocol represents a significant advancement in seed PA quantification. Its high-throughput capability, cost-effectiveness, and reproducibility make it an invaluable tool for screening large populations in breeding programs aimed at developing low-PA crop varieties. The successful identification of novel low-PA accessions highlights the potential of this method for accelerating crop improvement efforts focused on enhancing the nutritional quality of legumes and other crops.

## Acknowledgments

The author thanks Janneke Balk and Claire Domoney at the John Innes Centre for their guidance and support during this study.

